## Supplemental Data for "Group B *Streptococcus* Cas9 variants provide insight into programmable gene repression and CRISPR-Cas transcriptional effects"

Supplemental Figures

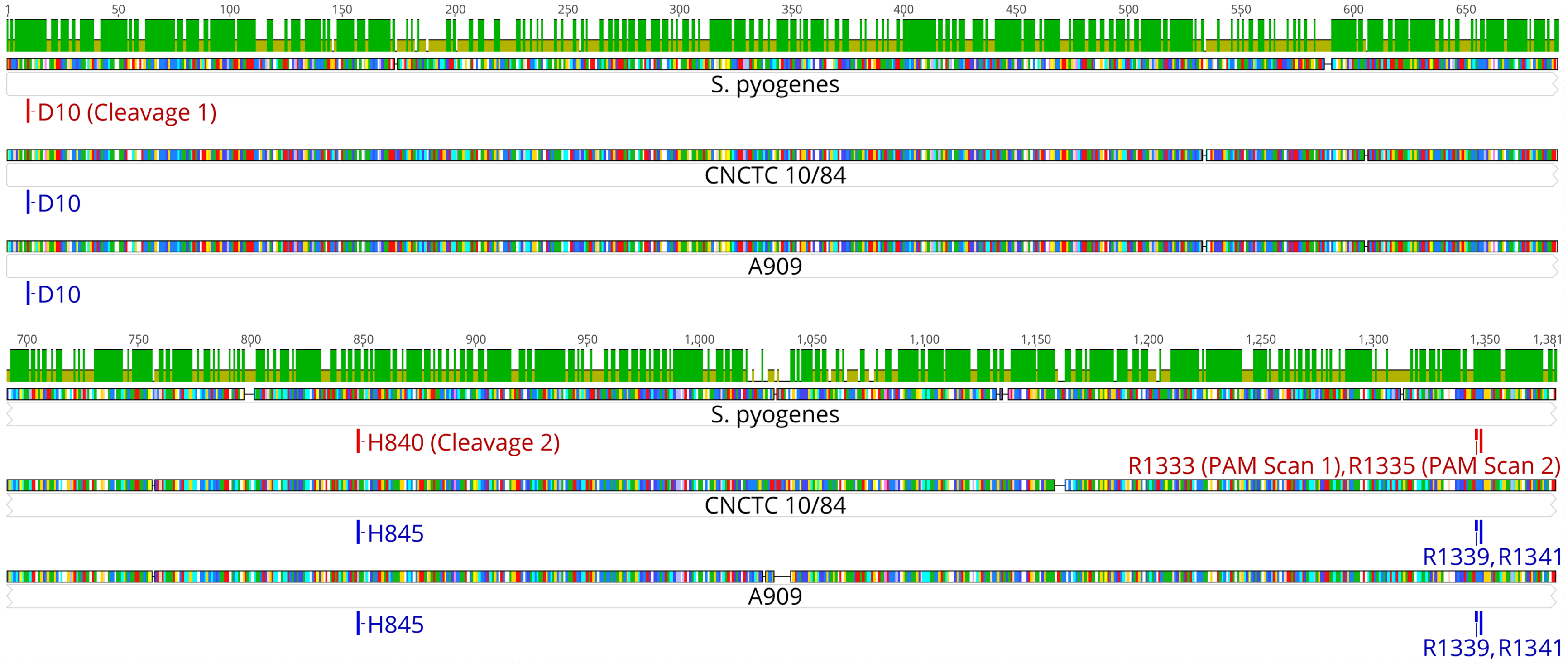


*Supplemental Figure 1*: Amino acid sequence alignments between group A *Streptococcus* (GAS; *S. pyogenes*) and two GBS strain Cas9 proteins. The top graph shows percent identity between the three proteins. The four active sites mutated in this study (two DNA cleavage sites and two PAM scanning sites) are indicated.

GAS* TRACR
AGCAAGUUAAAAUAAGGCUAGUCCGUUAUCAACUUGAAAAAGUGGCACCGAGUCGGUGC
UUUUUUU

GAS CR
GUUUUAGAGCUAUGCUGUUUUGAAUGGUCCCAAAAC

GAS sgRNA
$(N)_{20}$GUUUUAGAGCUAGAAAUAGCAAGUUAAAAUAAGGCUAGUCCGUUAUCAACUUGAAA
AAGUGGCACCGAGUCGGUGCUUUU

GBS* TRACR
AGCACAGCACGUUAAAAUAAGGCAGUGAUUUUUAAUCCAGUCCGUAUUCAGCUUGAAAA
AGUGAGCACCGAAUCGGUGCUUUUUUU

GBS CR
GUUUUAGAGCUGUGCUGUUUCGAAUGGUUCCAAAAC

GBS sgRNA
$(N)_{20}$GUUUUAGAGCUGUGAAACAGCACAGCACGUUAAAAUAAGGCAGUGAUUUUUAAUCC
AGUCCGUAUUCAGCUUGAAAAAGUGAGCACCGAAUCGGUGCUUUUUUU

* GAS sequences from M1 (NC_002737); GBS sequences from CNCTC 10/84

Predicted sgRNA folding (mFold)

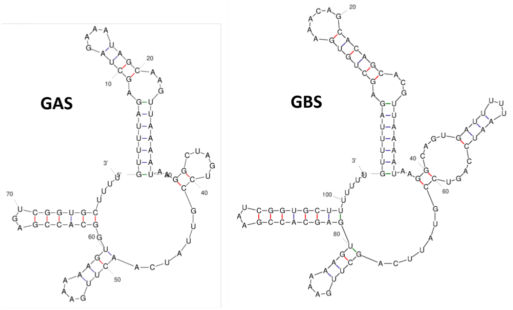


p3015b BsaI Cloning Site

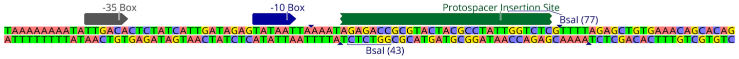


Protospacer cassette

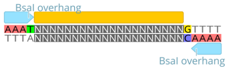


*Supplemental Figure 2*: **CRISPR-Cas nucleotide sequences from group A and group B *Streptococci*.** Orthologous RNA sequences are listed from *S. pyogenes* (GAS) and GBS, with red and blue highlighting to illustrate TRACR and CR conservation in sgRNA scaffolds derived from both strains. mFold analysis [101] shows expected stem-loop folding of sgRNA scaffolds. The cloning site of p3015b permits efficient insertion of a targeting protospacer cassette with phosphorylated termini complementary to the single strand overhangs left by BsaI linearization.

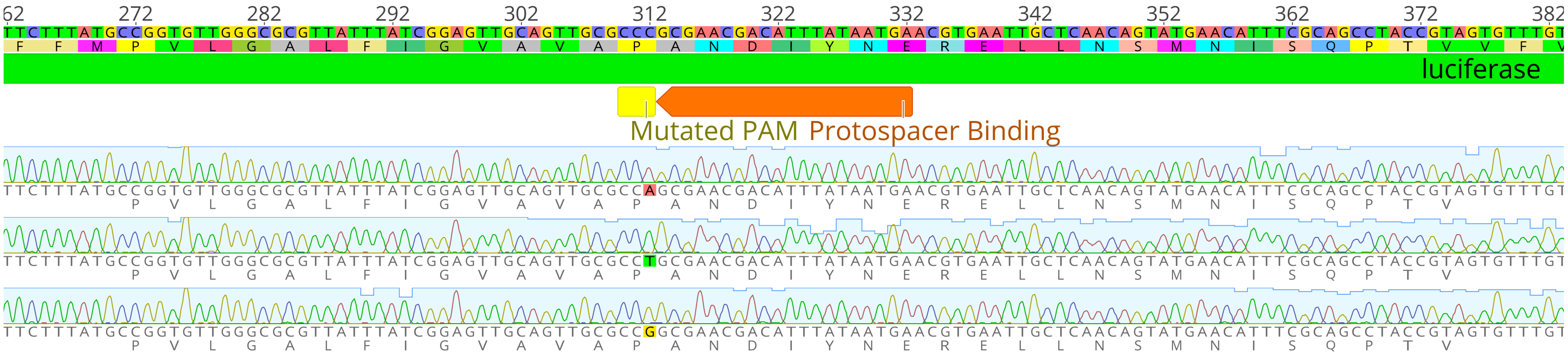


*Supplemental Figure 3*: **Sanger sequencing chromatograms for silent mutations of the CCC PAM in pFfluc to generate all four possible NGG variants.** The unmodified nucleotide sequence and its amino acid translation are illustrated at the top of the figure. For each chromatogram, the amino acid translation of the zoomed region is provided below the tracing.

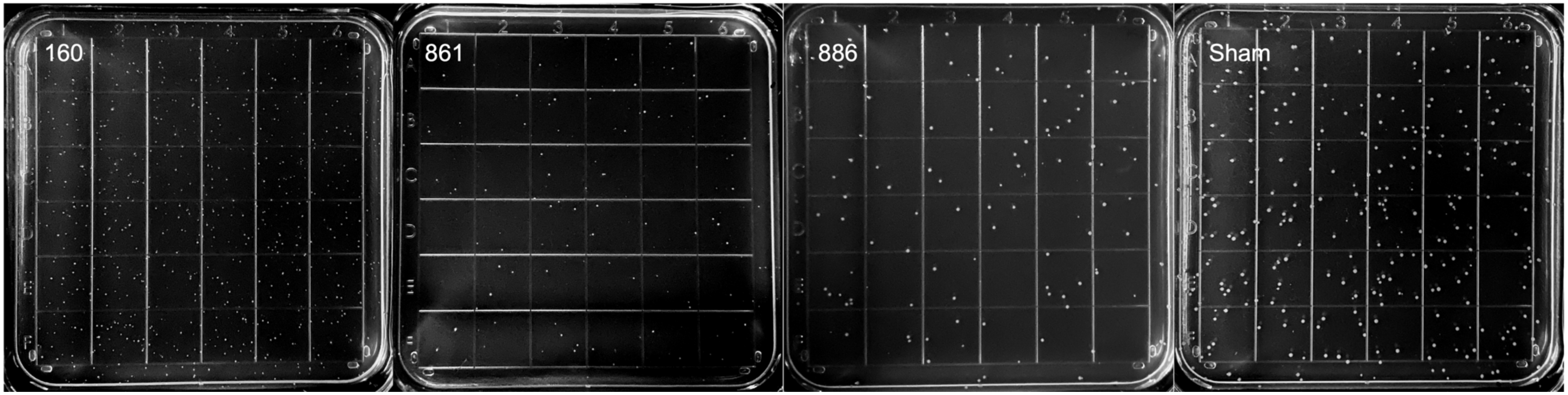


*Supplemental Figure 4*: **CRISPRi of the A909 *ccpA* gene, grown on tryptic soy plates, leads to colony sizes that correlate with growth kinetics in liquid cultures and reflect targeting site position within the gene.** Plates were photographed after 24-hours of growth following plating of O.D. normalized stock-cultures. Numbers above each plate indicate where along the *ccpA* coding sequence the protospacer targeted. This experiment was repeated three times. Representative photographs from one of the replicates are presented.

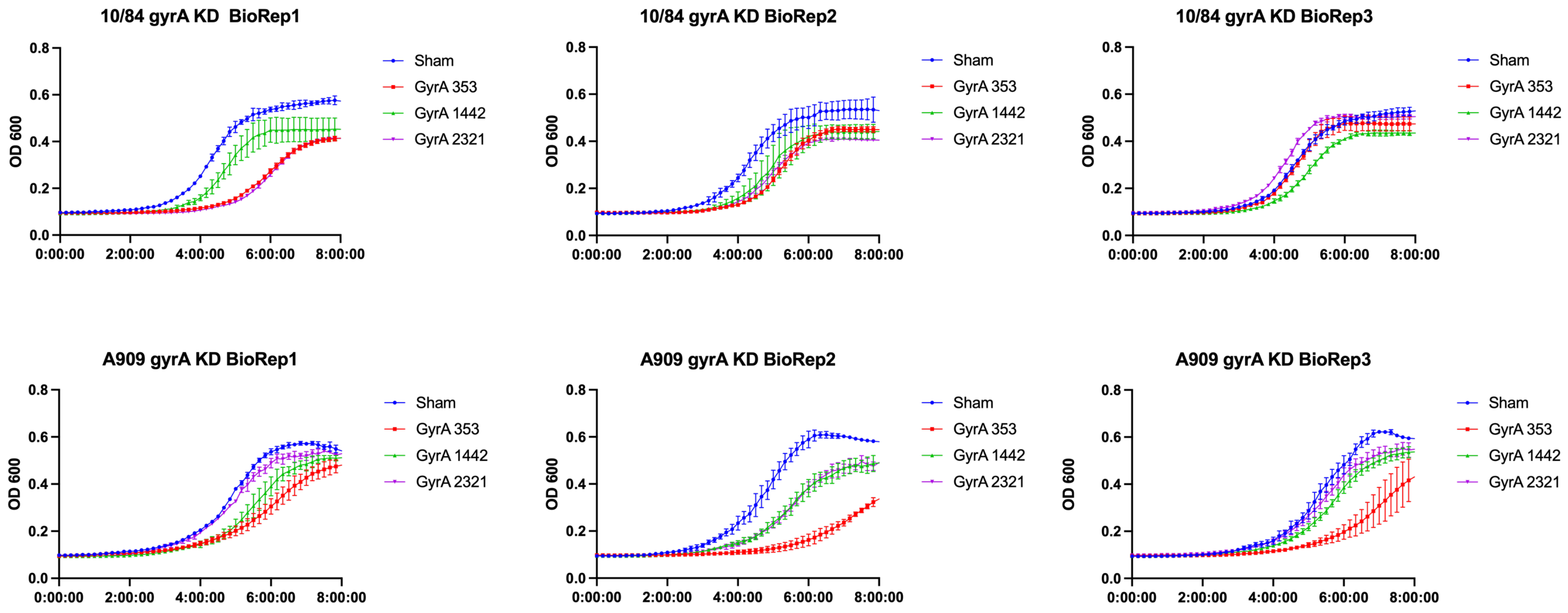


*Supplemental Figure 5*: **CRISPRi of the essential gene *gyrA* results in growth defects.** CRISPRi of *gyrA* in CNCTC 10/84 and A909 resulted in growth defects, with increased growth delay when the position of the targeted sequence was nearer to the 5’ end of the coding sequence. The numbers in the legend indicate the position of the protospacer along the *gyrA* gene. The experiment was conducted in three biological replicates, each with three technical replicates. The curves shown indicate the mean of all technical replicates at each time point. Error bars show standard deviation.

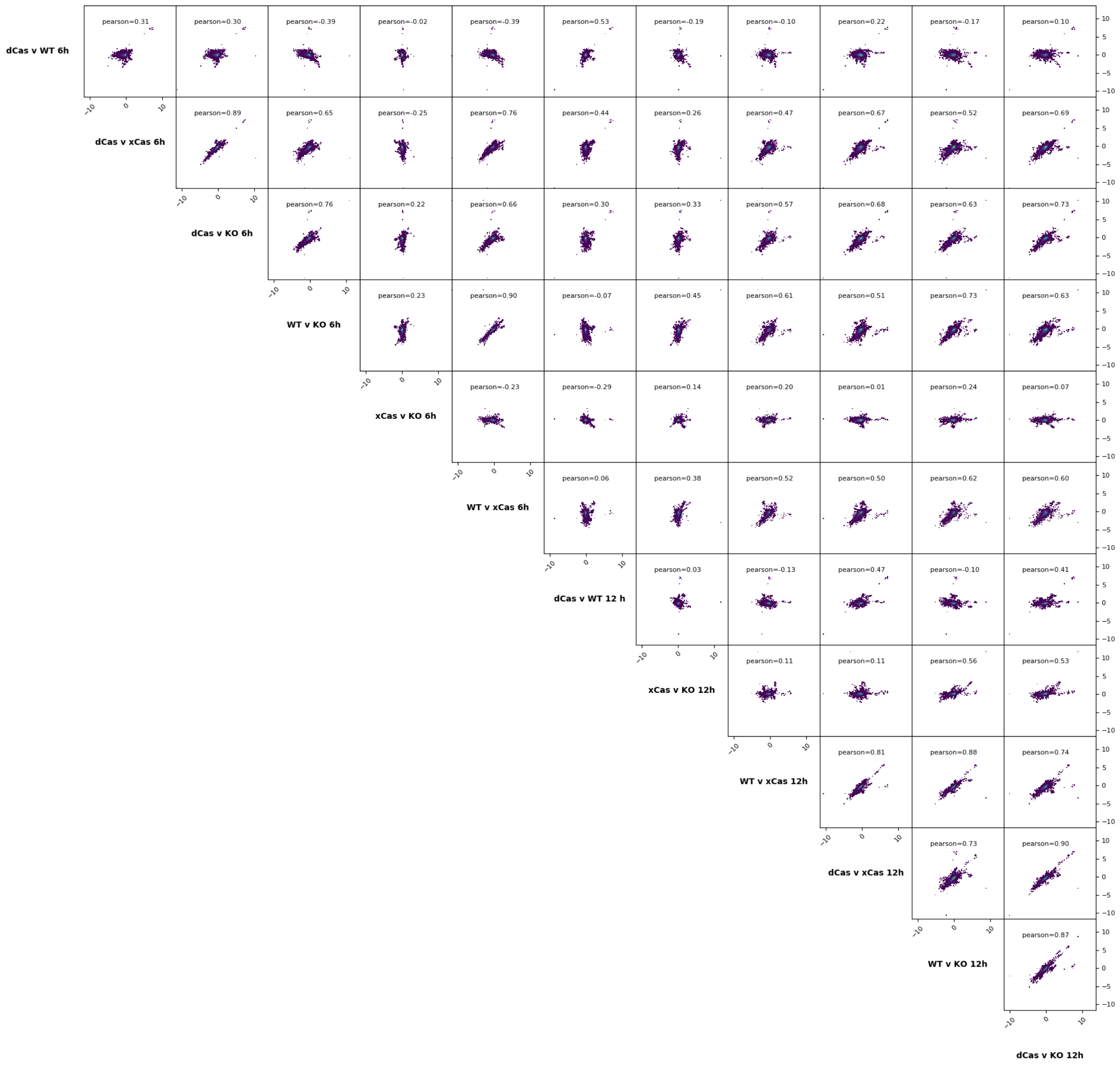


*Supplemental Figure 6*: **Pearson correlation scatterplots of CNCTC 10/84 Cas9 variant RNA-seq whole-genome expression data.**

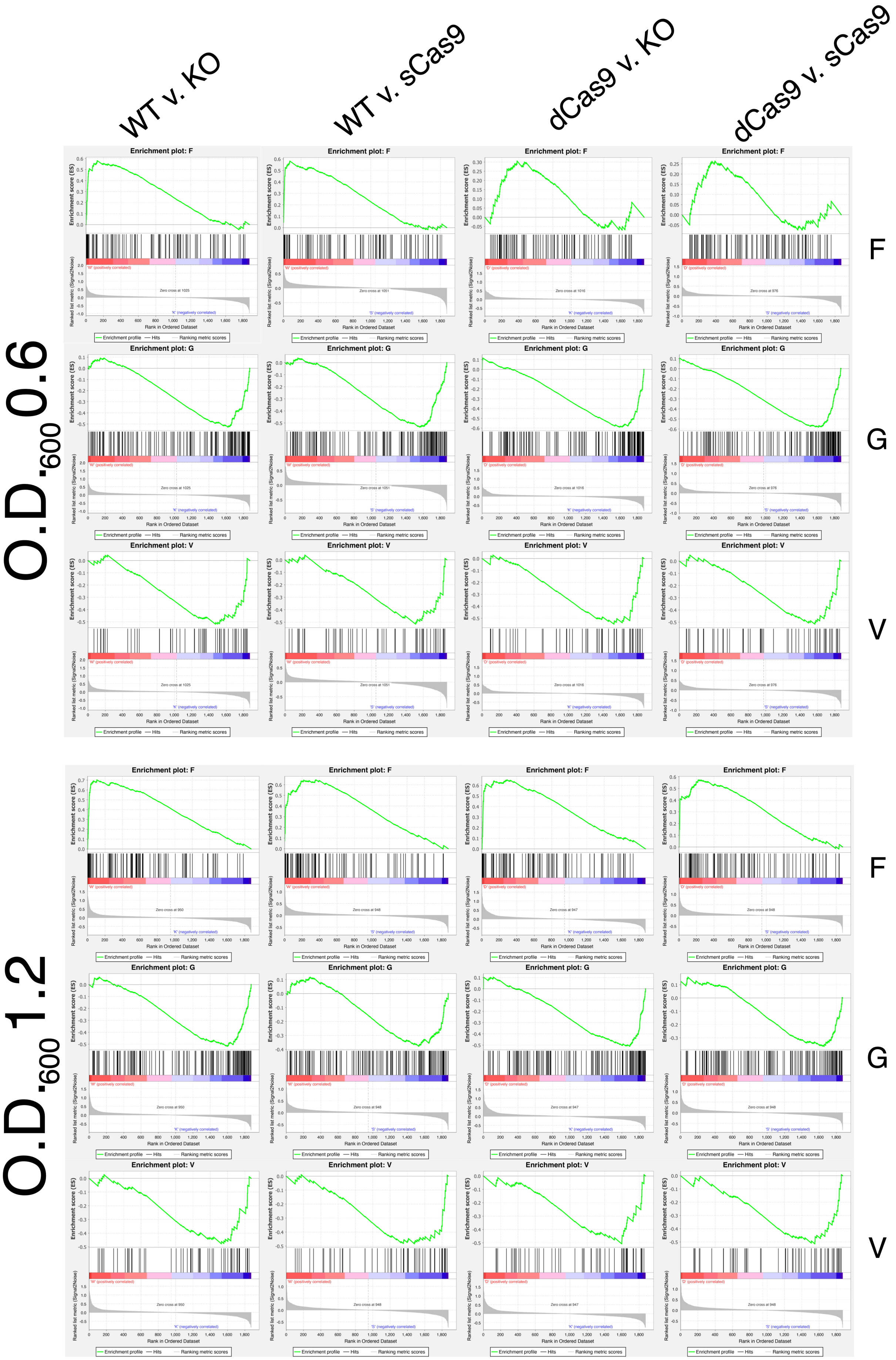

WT v. KO
WT v. sCas9
dCas9 v. KO
dCas9 v. sCas9
O.D.600 0.6
O.D.600 1.2
F
G
V
Enrichment plot: F
Enrichment plot: G
Enrichment plot: V
Enrichment score (ES)
Ranked list metric (Signal2Noise)
Rank in Ordered Dataset
Enrichment profile
Hits
Ranking metric scores

*Supplemental Figure 7*: **GSEA plots of GBS COG categories significantly enriched or depleted in PAM scanning v. non-scanning RNA-seq comparisons.** GSEA software generates a ranked list of all enriched or depleted genes, genome-wide, then assesses the distribution of genes within a functional category (for example, COG) within that list. Clustering of within-category genes at one end of the complete list or the other is quantified as an enrichment score, which is then normalized based on the size of the COG category [74]. These charts show the enrichment score graphs for genes within significantly enriched or depleted CNCTC 10/84 COG categories when comparing PAM scanning v. non-scanning RNA-seq differential expression data (see main text).

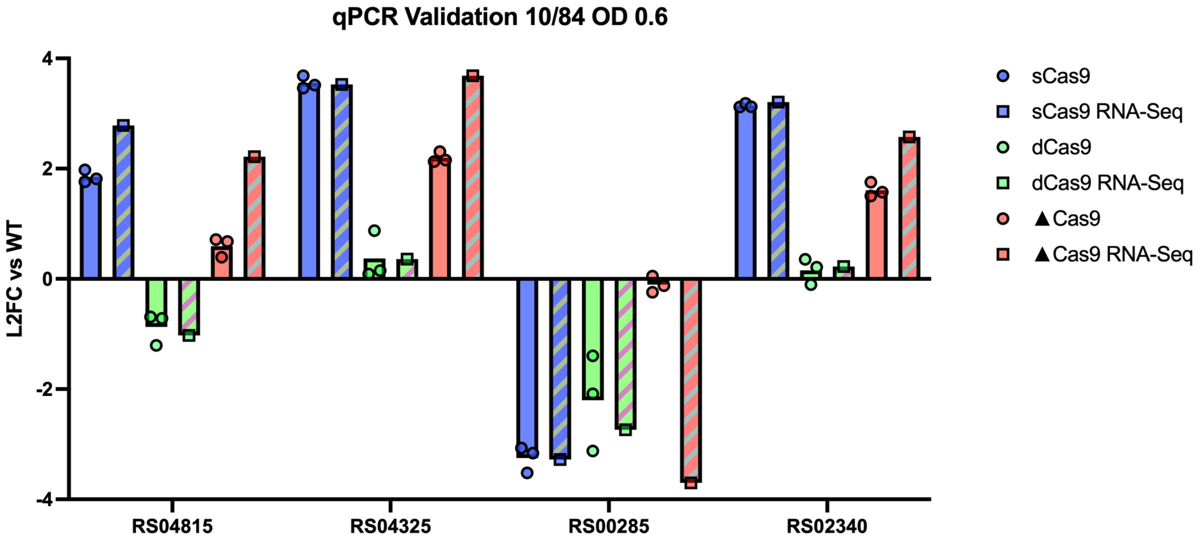


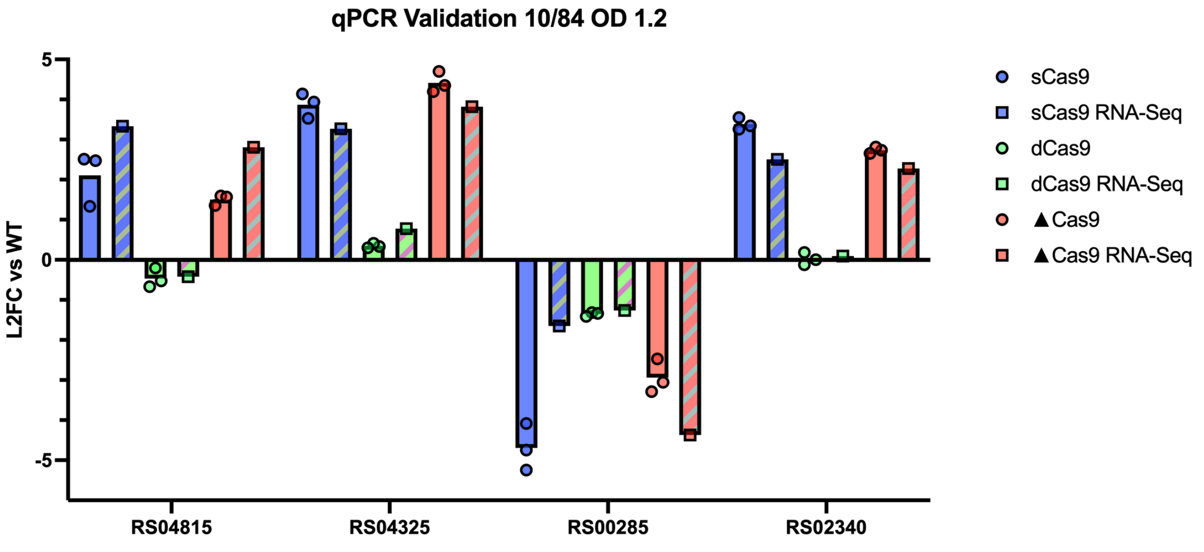


*Supplemental Figure 8*: **qRT-PCR validation of RNA-seq results.** cDNA from total RNA extracted from CNCTC 10/84 Cas9 variants was tested by qRT-PCR. Gene targets were selected for their consistent up- or downregulation across growth phases in RNA-seq data. The horizontal axis is labeled with RefSeq gene locus identifiers for the genes tested. Each sample was run in triplicate, along with reverse transcriptase-negative and no-template ($H_2O$ only) negative controls. Data from all replicates is shown. Comparisons to WT control cDNA were calculated, using *recA* expression as a normalization standard. $Log_2$ fold-change (L2FC) qRT-PCR results are plotted with identical comparisons from merged analyses of triplicate RNA-seq data. Bars indicate mean values, with error bars showing standard error of the mean for the qRT-PCR data.

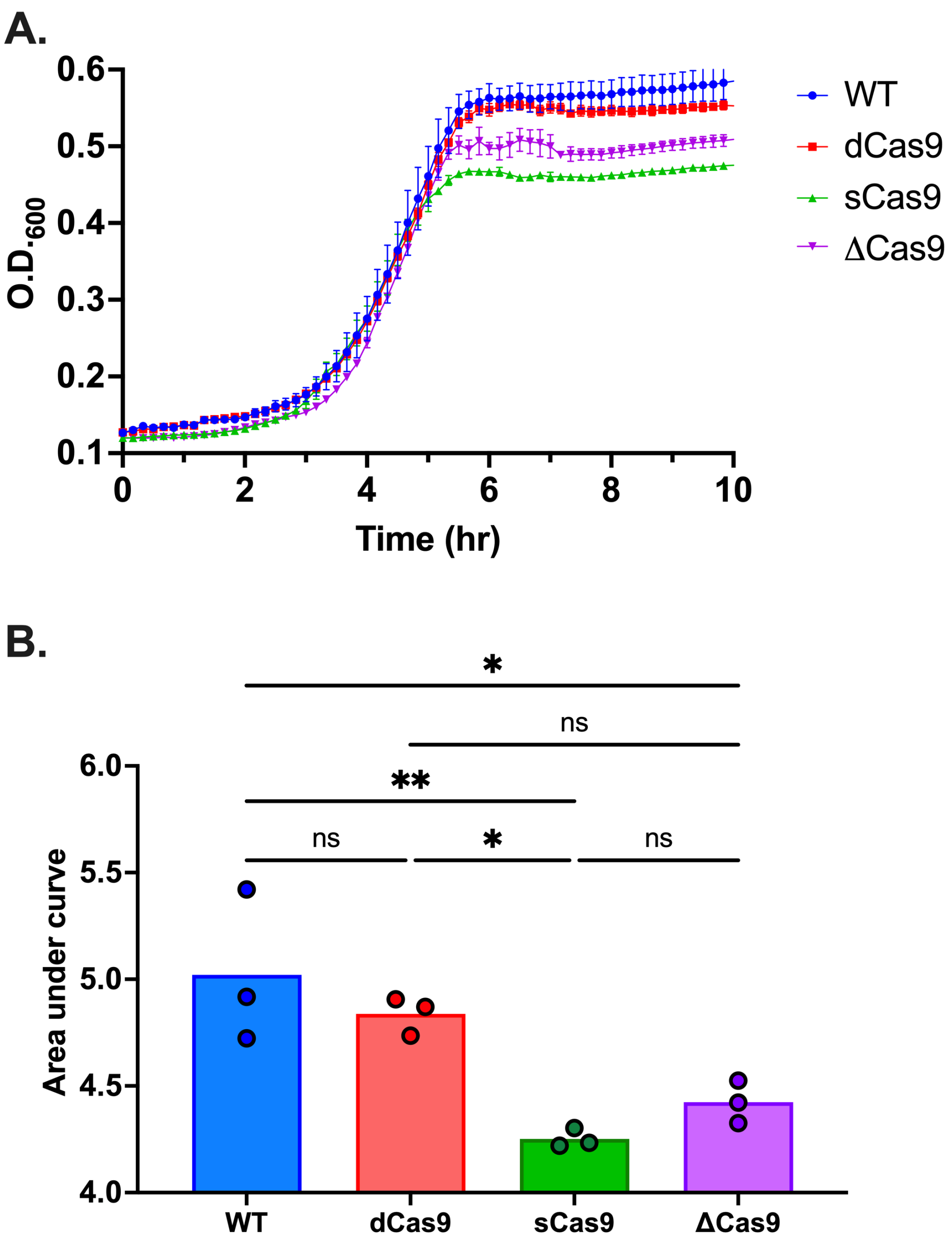


*Supplemental Figure 9*: In an *in vitro* oxidative stress challenge, CNCTC 10/84 *cas9* variants were grown with intermittent O.D.$_{600}$ monitoring (as described in Methods) in the presence of 5 mM porcine hemin (Fisher Scientific cat. #AAA1116503) added to liquid tryptic soy medium (**A**, n=3 biological replicates, curves show mean O.D.$_{600}$ with standard error of the mean for all replicates).

The experiment was conducted three times with separate biological replicates for each variant, using two technical replicates in the second and third repeats. Data points show mean $OD_{600}$ readings across the three independent experiments. Error bars show standard deviation. Area under the curve for the three biological replicates was calculated and compared by one-way ANOVA with Bonferroni correction for multiple comparisons (**B**; * $p<0.05$, ** $p<0.01$, *** $p<0.005$, **** $p<0.001$).

Supplemental Data

**Supplemental Data 1:** Plasmids and primers used in this work.

| Plasmid | Description | Antibiotic resistance | Reference |
|---|---|---|---|
| pMBsacB | Temperature- and sucrose-sensitive broad host-range mutagenesis plasmid | Erythromycin | [90] |
| p3015b | Broad host-range shuttle vector with GBS-compatible sgRNA expression cassette | Erythromycin | From the pVPL3004 backbone in Oh J and van Pijkeren, 2014[17] |
| pFfluc | Firefly luciferase expression vector | Kanamycin | From pFfluc-Erm with antibiotic resistance cassette changed to kanamycin[79] |
| pDC123 | Broad host-range plasmid used for chloramphenicol resistance gene amplification | Chloramphenicol | [92] |

| Primer name | Sequence | Description |
|---|---|---|
| dcas9_mut1_F | GATTAAAAATCACTGCCTTATTTTAACGTGCTGTGC | Amplification of *dcas9* D10A fragment from custom synthesized DNA |
| dcas9_mut1_R | TGGAGCTCCACCGCGGTG | Amplification of *dcas9* D10A fragment from custom synthesized DNA |
| dcas9_mut2 _F | AAAAACAATCTTGGACTATCTTATTG | Amplification of *dcas9* H845A fragment from custom synthesized DNA |
| dcas9_mut2 _R | TGGCTCTAACTGAGGATATTTG | Amplification of *dcas9* H845A fragment from custom synthesized DNA |
| scas9_US_F | TCTAAGGAATCAATCTTGGCG | Amplification of *scas9* fragment 1 |
| scas9_US_R | GATGTATAGGCCATCAACTATTTTTGTTTATC | Amplification of *scas9* fragment 1 with R1339A |

| | | |
|---|---|---|
| | | and R1441A mutations at the 3' terminus |
| scas9_DS_F | GTTGATGCAAAAGCCTATACATCAACTAAAG | Amplification of *scas9* fragment 2 with R1339A and R1441A mutations at the 5' terminus |
| scas9_DS_R | GGTTCTAAGTCATGGTAGAGATTC | Amplification of *scas9* fragment 2 |
| xcas9_US_F | TCCAGGAGATTCTACTTG | Amplification of *cas9* upstream fragment for assembly of Δ*cas9* mutagenesis cassette |
| xcas9_US_R | AACTTTTCTCCTTTTATTTAAAACAC | Amplification of *cas9* upstream fragment for assembly of Δ*cas9* mutagenesis cassette |
| xcas9_DS_F | TATGGCAGGTTGGCGAAC | Amplification of *cas9* downstream fragment for assembly of Δ*cas9* mutagenesis cassette |
| xcas9_DS_R | TTGAAAAATCATTTCCAAAAAGTGTATTG | Amplification of *cas9* downstream fragment for assembly of Δ*cas9* mutagenesis cassette |
| xcas9_cat_F | ATGAACTTTAATAAAATTGATTTAGACAATTG | Amplification of *cat* chloramphenicol resistance gene for assembly of Δ*cas9* mutagenesis cassette |
| xcas9_cat_R | TTATAAAAGCCAGTCATTAGG | Amplification of *cat* chloramphenicol resistance gene for assembly of Δ*cas9* mutagenesis cassette |

**Supplemental Data 2**: Protospacer sequences used in this work. For cloning into p3015b, each forward protospacer is preceded by AAAT at the 5' end and followed by a G at the 3' end. The reverse protospacer is preceded by AAAAC at the 5' end. These additional bases confer complementarity to the BsaI overhangs in the linearized plasmid.

| **Gene target** | **Predicted cut position** | **Forward protospacer*** | **Reverse protospacer*** |
|---|---|---|---|
| Sham | N/A | GTACTACGCCTATTGGTCTC | GAGACCAATAGGCGTAGTAC |
| *gyrA* | 353 | TGCTTCTGTATAACGCTGTG | CACAGCGTTATACAGAAGCA |
| *gyrA* | 1442 | AATCATTAATTCTGTACGAC | GTCGTACAGAATTAATGATT |
| *gyrA* | 2321 | TTGAGAAATATTAGCAACAT | ATGTTGCTAATATTTCTCAA |
| *ccpA* | 160 | AAACCACGCGCCACAGCATT | AATGCTGTGGCGCGTGGTTT |
| *ccpA* | 861 | TGACTGATAGAAGTTAAGTT | AACTTAACTTCTATCAGTCA |
| *ccpA* | 886 | ACAGCACCCAAATCATAGAC | GTCTATGATTTGGGTGCTGT |
| *ffluc* | 316 | TCATTATAAATGTCGTTCGC | GCGAACGACATTTATAATGA |
| *ffluc* (+) | 267 | CTCTCTTCAATTCTTTATGC | GCATAAAGAATTGAAGAGAG |
| *ffluc* | 1189 | ACGCCCGCGTCGAAGATGTT | AACATCTTCGACGCGGGCGT |
| *ffluc* | 1390 | ACATAACCGGACATAATCAT | ATGATTATGTCCGGTTATGT |
| *ffluc* | -34 | TGAATGACAATGATGTTCCG | CGGAACATCATTGTCATTCA |
| *ffluc* | -76 | TAGAATTAAATTAAAAAGGG | CCCTTTTTAATTTAATTCTA |
| *ffluc* | 68 | CTTATGCAGTTGCTCTCCAG | CTGGAGAGCAACTGCATAAG |
| *ffluc* | 277 | ATAAATAACGCGCCCAACAC | GTGTTGGGCGCGTTATTTAT |
| *cyl* operon | 283 | ATCCTTTCCATTAATAGTAT | ATACTATTAATGGAAAGGAT |
| *cyl* operon | 384 | GAAAAAAGAATTTCACCTTT | AAAGGTGAAATTCTTTTTTC |
| *covR* | 227 | TGCTGTCATCATCATGATAT | ATATCATGATGATGACAGCA |
| *covR* | 313 | AATAATTCTTCGATTGCAAA | TTTGCAATCGAAGAATTATT |

*Predicted cut position refers to the predicted site that WT Cas9 would cleave upon target identification. It is relative to the 5' start of the target gene or operon. The corresponding bases in the forward and reverse protospacer sequences are indicated in red.

**Supplemental Data 9:** Primer sequences for qRT-PCR validation of RNA-seq results.

| | **qPCR Primers** | |
|---|---|---|
| **Genes** | **Forward** | **Reverse** |
| *cyl* | ACGTCGTGCCCAACTTATAG | ACACCAGCTCTATCATCAGTAATAA |
| *covR* | TGCGTGGTGATGAGGAAAT | AGCAACTCTTCTCGTGTCATAA |
| *ccpA* | GGCAGGATACAAAGAAGGTCTC | CACACGTTGAGCTAATGCAAAG |
| RS04815 | CACAGAGCGACACTATCTTTGA | TTGGTAAGATAAGCTCCGCTATT |
| RS04325 | GCAGAAGCGATTGCAAGTATG | CGCTGCTTCATCCACAATAATC |
| RS00285 | TGGAGCTCGTAAGGAGACTATT | GTCACCACACCAGCCATATT |
| RS02340 | CGGACCAGCTAAAGACAGATAG | GGCATCTGCACTAGCCTTATTA |
| *recA* | GTGGGATTGCTGCCTTTATTG | CTGAGTCAGGTTGAGACAAGAG |
